# Structural proteomics of a bacterial mega membrane protein complex: FtsH-HflK-HflC

**DOI:** 10.1101/2023.06.07.544024

**Authors:** Hatice Akkulak, H. Kerim İnce, Gunce Goc, Carlito B. Lebrilla, Burak V. Kabasakal, Sureyya Ozcan

## Abstract

Recent advances in mass spectrometry (MS) yielding sensitive and accurate measurements along with developments in software tools have enabled the characterization of complex systems routinely. Thus, structural proteomics and cross-linking mass spectrometry (XL-MS) have become a useful method for structural modeling of protein complexes. Here we utilized commonly used XL-MS software tools to elucidate the protein interactions within a membrane protein complex containing FtsH, HflK, and HflC, over-expressed in *E.coli*. The MS data were processed using MaxLynx, MeroX, MS Annika, xiSEARCH, and XlinkX software tools. The number of identified inter- and intra-protein cross-links varied among software. Each interaction was manually checked using the raw MS and MS/MS data and distance restraints to verify inter- and intra-protein cross-links. A total of 37 inter-protein and 148 intra-protein cross-links were determined in the FtsH-HflK-HflC complex. The 59 of them were new interactions on the lacking region of recently published structures. These newly identified interactions, when combined with molecular docking and structural modeling, present opportunities for further investigation. The results provide valuable information regarding the complex structure and function to decipher the intricate molecular mechanisms underlying the mega FtsH-HflK-HflC complex.

Graphical Abstract
The overall workflow of XL-MS analysis: the steps of XL-MS studies applied to the two samples taken from different stages of purification process of mega membrane protein complex FtsH-HflC-HflK.

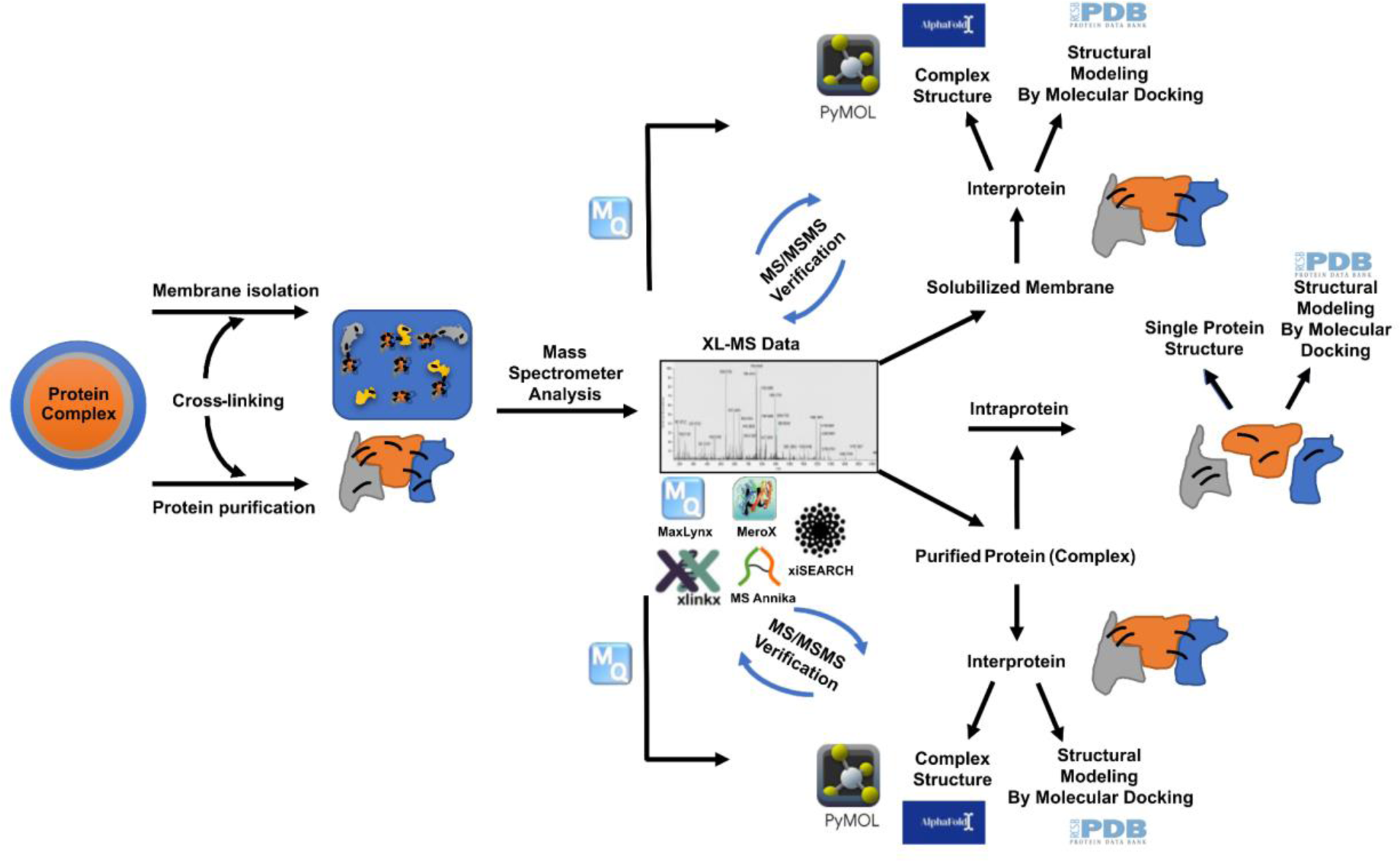

## 1. INTRODUCTION

The role and function of proteins in the cell could only be revealed by uncovering all aspects of a protein: structure, alterations, localizations, abundances, and the protein-protein interactions that are the basis of proteomics research. Since proteins might gain their function through forming complexes with various proteins under various conditions, functional proteomics is one of the most effective approaches to understand the role of the proteins by identifying protein-protein interactions [1].

Recent developments in proteomics with more rapid and sensitive mass spectrometry (MS) analyses have enabled us to determine various features in a single MS run [2]. Current software used in MS allows the analysis of thousands of spectra, leading to deciphering even very complex matrices to unravel thousands of features. Software tools have made MS spectra analysis processes faster and automated, which aided large-scale clinical and multidisciplinary studies [3].

Proteomics has become a fashionable method for structural studies rather than a complementary technique by providing comprehensive information especially in protein-protein interactions [4,5]. X-ray crystallography and electron cryo-microscopy (cryo-EM) often reveal only one form of a protein complex with interactions defined by the organization of the protein components whereas cross-linking mass spectrometry (XL-MS) can give overall information from multiple conformations and compositions of a protein complex simultaneously. XL-MS software are designed to identify interacting residue pairs and give information about connections between these residues by utilizing spectrum data. Numerous software [6] are available, including Kojak [7], MaxLynx [8], MeroX (StavroX) [9–12], MS Annika [13], OpenPepXL [14], pLink2 [15], XlinkX [16], and XiSEARCH [17], for various cross-linking applications.

Crucial processes of the cell, such as signal transduction, membrane transport, signal detection, cell-cell interactions are operated by membrane proteins. Therefore, 25% of cell proteomes are composed of membrane proteins, making them an important target for developing new therapeutics [18]. Folding and degradation of membrane proteins are controlled by chaperones and proteases [19]. As a member of AAA protease family, a conserved protein FtsH has a cytoplasmic ATPase domain and possesses a proteolytic activity with a zinc-binding site to degrade the improperly folded integral membrane proteins [20,21]. HflK and HflC are known to form a very large membrane-bound complex with FtsH, and modulate FtsH proteolytic activity [19,21,22]. FtsH-HflK-HflC complex plays a chaperone role to stabilize proteins in mitochondria [20], and a modulator role to limit ATPase activity of FtsH [22,23].

Here we used a cross-linking MS approach to investigate the structure and the residue-residue interactions of a recombinantly produced bacterial membrane protein complex composed of three proteins, FtsH, HflK, and HflC. We utilized commonly used XL-MS software, MaxLynx, MeroX, MS Annika, xiSEARCH and XlinkX for the analysis of both the purified protein complex and total solubilized membrane samples. Our findings will give structural insights into FtsH-HflK-HflC complex and guide structural proteomics studies using XL-MS.

## 2. MATERIALS AND METHOD

### 2.1. Expression and Purification of FtsH-HflK-HflC Membrane Protein Complex

The plasmid containing FtsH-HflK-HflC membrane protein complex genes was transformed into *E. coli* Lemo21 (DE3) (NEB, GB) for protein expression. Overexpression was carried out by using 0.4 mM IPTG (Sigma, USA) and 0.2% (w/v) arabinose (Carl Roth, DEU) in 2xYT media (Sigma, USA). 10 g of harvested cells were disrupted by sonication (Sonics VCX 130, USA). Afterwards, the membrane was separated by ultracentrifugation, and solubilized using 2% (w/w) n-Dodecyl-β-D-maltoside (DDM) (Carbosynth, USA) and purified samples of FtsH-HflK-HflC were collected using affinity chromatography.

### 2.2. Chemical Cross-Linking Reactions with DSBU and Tryptic Digestion

Cross-linking reactions with disuccinimidyl dibutyric urea (DSBU) (Thermo Fisher Sigma, USA) were carried out using purified FtsH-HflK-HflC membrane protein complex and solubilized membrane with overexpressed protein complex according to manufacturer’s instructions [11,24–26]. A 5 µM of purified FtsH-HflK-HflC sample in 50 mM HEPES-NaOH pH 8.0, 150 mM NaCl buffer was treated with 2.5 mM DSBU for 60 min at room temperature. The reaction was stopped using 20-fold Tris-HCl (1 M, pH 8.0) and samples were concentrated using pre-equilibrated 100 kDa MWCO concentrators up to ∼20 µL.

In addition to the purified FtsH-HflK-HflC complex, cross-linking reactions were also performed for solubilized membranes of FtsH-HflK-HflC. Reactions were set up at 100 mg/ml solubilized membranes with a final concentration of 10 mM DSBU in 12.5 µL. Reactions were incubated for 60 min at room temperature and terminated using 20-fold Tris-HCl (1 M, pH 8.0).

Samples were extracted in 100 µL of 50 mM Ammonium bicarbonate and incubated with 11 µL of 100 mM Dithiothreitol (DTT) for 50 min at 60 °C. Free cysteines were alkylated with 22 µL of 100 mM Iodoacetamide (IAA) for 30 min at room temperature in the dark. Then 50 µL 0.1 µg/µL trypsin was added and samples were incubated at 37 °C for 18 h. All samples were dried in miVac at room temperature. Dry peptide samples were dissolved in 0.1% Formic acid and peptide concentrations were determined with Pierce™ Quantitative Peptide Assay (Thermo Fisher Scientific, USA) before the MS analysis.

### 2.3. Mass Spectrometry

The 1 µg digested samples were analyzed by the UltiMate™ WPS-3000RS nanoLC system coupled with Orbitrap Fusion Lumos (Thermo Fisher Scientific, USA). The peptides were separated on Acclaim^TM^ PepMap^TM^ 100 C18 HPLC Columns (3 μm, 0.075 mm × 500 mm, Thermo Fisher Scientific, USA). The mobile phase A containing 0.1% aqueous formic acid and mobile phase B comprising 0.1% formic acid in 80% acetonitrile were set to a gradient; 0–5 min, 4–4% (B); 5–130 min, 4–35% (B); 130–150 min, 35–50% (B); 150–153 min, 50–100% (B); 153–168 min, 100–100% (B); 168–170 min, 100–4% (B); and 170–180 min, 4–4% (B). The MS and MSMS spectra were collected with a mass range of m/z 300-1800 in positive ionization ion mode. The high-energy C-trap dissociation (HCD) fragmentation was performed with nitrogen gas, and collision energies of 25%, 30%, and 35%. The precursor and the product ions were detected at 120K resolution and 15K resolution, respectively.

### 2.4. Data Analysis

Raw MS data corresponding to the purified protein complex and the solubilized membrane were analyzed in MaxLynx, MeroX 2.0, MS Annika, xiSEARCH and XlinkX to identify cross-linked peptides as described below.

MaxLynx software is an add-on to the MaxQuant. MS Annika and XlinkX are two separate nodes for the Proteome Discoverer 2.5 (Thermo Fisher Scientific). Corresponding workflows were obtained from the website. MeroX and xiSEARCH are the JAVA-based software that can be operated independently.

Protein search was carried out through *E.coli* proteome (UniProt ID: UP000000625) by MaxQuant to identify the most abundant proteins. Cross-link search was carried out with three complex proteins FtsH-HflC-HflK (UniProt IDs: P0AAI3-P0ABC3-P0ABC7) sequences for purified protein complex sample. The top 300 proteins from protein search results from MaxQuant, and the MaxQuant contaminants file (Supplementary Table 1) were combined to create a joint FASTA file for the search of solubilized membrane sample [27]. While the raw MS data could be directly used in MaxLynx, MS Annika and XlinkX, it needs to be converted to a mzml format using an external software such as ProteoWizard [28] for MeroX, and mgf format for xiSEARCH. The same settings were used in all five software to conduct searches under the same conditions (Supplementary Table 1). C-terminals of lysine (K) and arginine (R) residues were set as specific cleavage sites, and maximum missed cleavages were set to 3. Precursor and fragmentation mass tolerance were limited to 10 ppm and 20 ppm, respectively. The carbamidomethylation at cysteine was assigned as the fixed modification, and the oxidation at methionine was selected as the variable modification. DSBU and its modifications were introduced as chemical modifications. Signal to noise ratio was set to 2.0. The minimum precursor mass limit was 1000 Da, whereas the maximum precursor mass limit was 20,000 Da. In addition, the minimum and maximum peptide lengths were set as 5 and 40 AAs. The cross-link modification sites were specified as Lysine for all software. In addition, Serine (S), Threonine (T), and Tyrosine (Y) were also added in MeroX and xiSEARCH search as the software enables variable amino acid search [29]. The FDR (False Discovery Rate) was limited to 1%. The upper limit to Cα-Cα distance was used as 30Å as suggested by previous studies [12,30,31].

### 3. RESULTS AND DISCUSSION

This study employs a software-based approach to get insights into a large membrane protein complex structure using XL-MS. The intra- and inter-protein cross-links within and between the large multimeric membrane protein complex, FtsH-HflK-HflC were analyzed in the purified form and over-expressed in the solubilized membrane to obtain structural information. The proteins were cross-linked with DSBU and then digested by using trypsin. High-resolution MS spectra were analyzed using common XL-MS software MaxLynx, MeroX, MS Annika, xiSEARCH, and XlinkX. The intra-protein and inter-protein cross-links were further confirmed using MS and MS/MS data and distance restraints.

### 3.1. XL-MS Analysis

To identify inter- and intra-protein cross-links, signature fragment ions representing cross-linkers and peptides were searched through five software.

A representative MS/MS spectrum of a cross-linked peptide pair is presented in Figure 1. The MS cleavable cross-linker, DSBU, provided a significant advantage over non-cleavable cross-linkers, as software were able to confirm specific spectral assignments using the mass difference between two cleavages of cross-links with the backbone fragmentation products. Moreover, cleavable cross-linkers provide more sequence coverage with backbone fragmentation of peptides [32]. The DSBU was covalently connected to residues of proteins using NHS ester reactive sides. The Higher-energy collisional dissociation (HCD) cleaves DSBU from amide bonds and results in two characteristic diagnostic ions. The diagnostic ion corresponds to the peptide and cross-linker with protonated amine group at the cleavage point (PEPTIDE ½+Bu), which provides 85u mass adduct on the peptide [11,12]. In Fig. 1, the FGKVLR peptide with Bu part of the cross-linker on the lysine gives the peak of 803.5043. Also, the VTAETKGK peptide with Bu part of the cross-linker on the lysine peak appears at 917.5200. The other diagnostic ion is the BuUr part of the cross-linker bound to the peptide, where the nitrogen on amine group loses one proton due to the cross-linker cleavage, resulting in a 111u mass adduct to the connecting peptide (PEPTIDE ½+BuUr). Fig. 1 shows the FGKVLR peptide with BuUr part of the cross-linker on the lysine giving the peak of 829.4832, and the VTAETKGK peptide with BuUr part of the cross-linker on the lysine peak appearing at 943.5001. The cross-link search was conducted using specific software designed to search 26u mass difference owing to the cleavage of the cross-linker described above [8,12,13,16].

**Figure 1.**
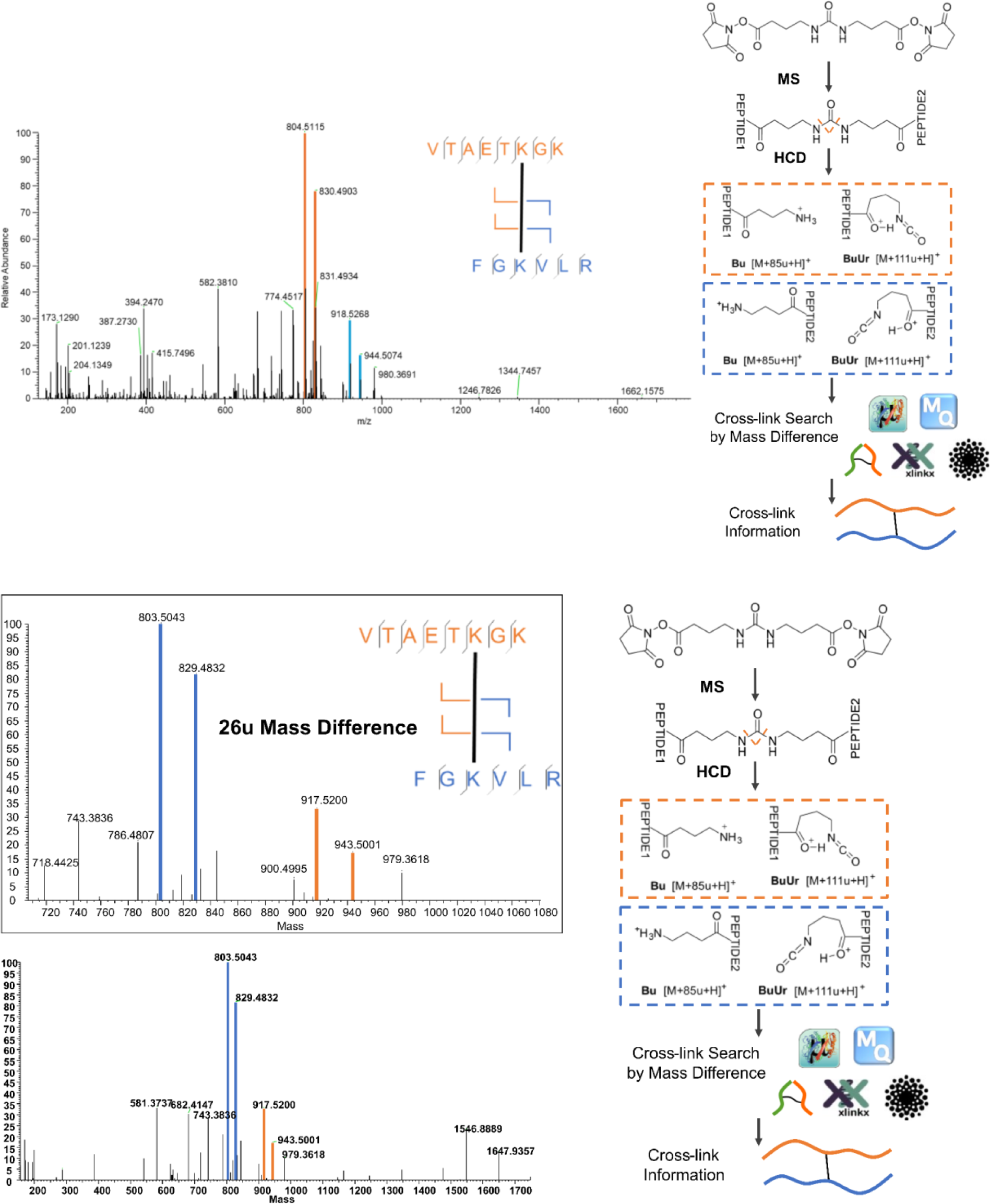
Fingerprint fragments confirming cross-linked peptides.

The XL-MS software predominantly uses lysine-specific cross-linking sites to identify cross-links. While the literature suggests that serine, threonine, and tyrosine could also be cross-linked to DSBU [29], it’s often noted that most of these S, T, Y cross-links are false positives. The simultaneous multi/variable amino acid-cross-linker searches by MeroX and xiSEARCH enabled us to identify the DSBU connection other than lysine. A representative fragmentation pattern confirming threonine-DSBU connection is shown in Fig. 2.

**Figure 2.**
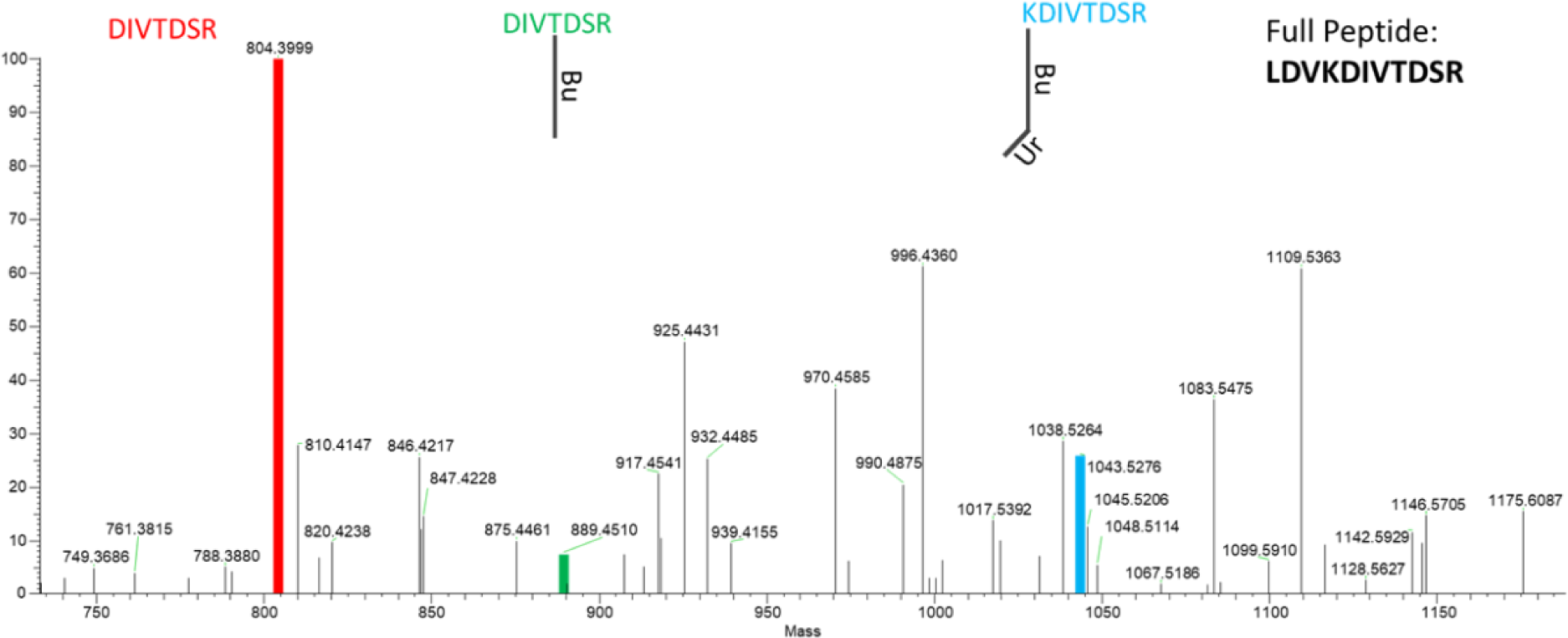
Different cleavage patterns of the cross-linker and backbone revealing the cross-link between threonine and lysine.

The deconvoluted MS/MS spectrum of LDVKDIVTDSR-GVIGKYTmDR peptide pair is given in Fig. 2. In addition to two diagnostic ions specific to peptide-DSBU (85 and 111), several ions were observed to validate the threonine-DSBU connection. The peak at the 1043.5276 mass unit belongs to Y8 cleavage of the peptide LDVKDIVTDSR and BuUr part of the cross-linker which is KDIVTDSR+BuUr, suggesting the cross-linker is bound to lysine. The peak at 889.4510 mass unit belongs to Y7 cleavage of the peptide LDVKDIVTDSR and Bu part of the cross-linker which is DIVTDSR+Bu. In this example, there is no lysine in the sequence, but threonine. Furthermore, the peak at 804.3999 mass unit belongs to Y7 cleavage of the LDVKDIVTDSR yielding the DIVTDSR without the cross-linker. Data suggested that the DSBU was linked to both lysine and threonine residues and corresponding peptides were co-eluted.

The cross-link search through five XL-MS software from two experimental setup yielded 185 unique cross-links (Supplementary Table 2-1). The total number of inter-protein and intra-protein connections were 37 and 148, respectively.

### 3.2. FtsH-HflK-HflC Complex Inter-protein Cross-links

A total of 37 unique inter-protein cross-links were obtained from the two samples (Supplementary Table 2-2). Among them, one key interaction between FtsH and HflK was only found by MaxLynx from the purified protein sample. The number of cross-links between FtsH and HflC were 14, whereas the number of cross-links between HflK and HflC was 22 from all the software and samples. Five of the cross-links were found through all five software considering both samples. The highest number of unique cross-links was obtained from the purified protein complex sample by MS Annika. In total, the least number of unique cross-links was found in the solubilized membrane sample.

There were three common inter-protein cross-links obtained from purified protein samples and 2 from solubilized membrane samples with five software (Figure 3). All five software found individual cross-links not found by other software in the purified protein sample (Fig. 3a), whereas three software provided no individual cross-links in the solubilized membrane sample (Fig. 3b).

**Figure 3.**
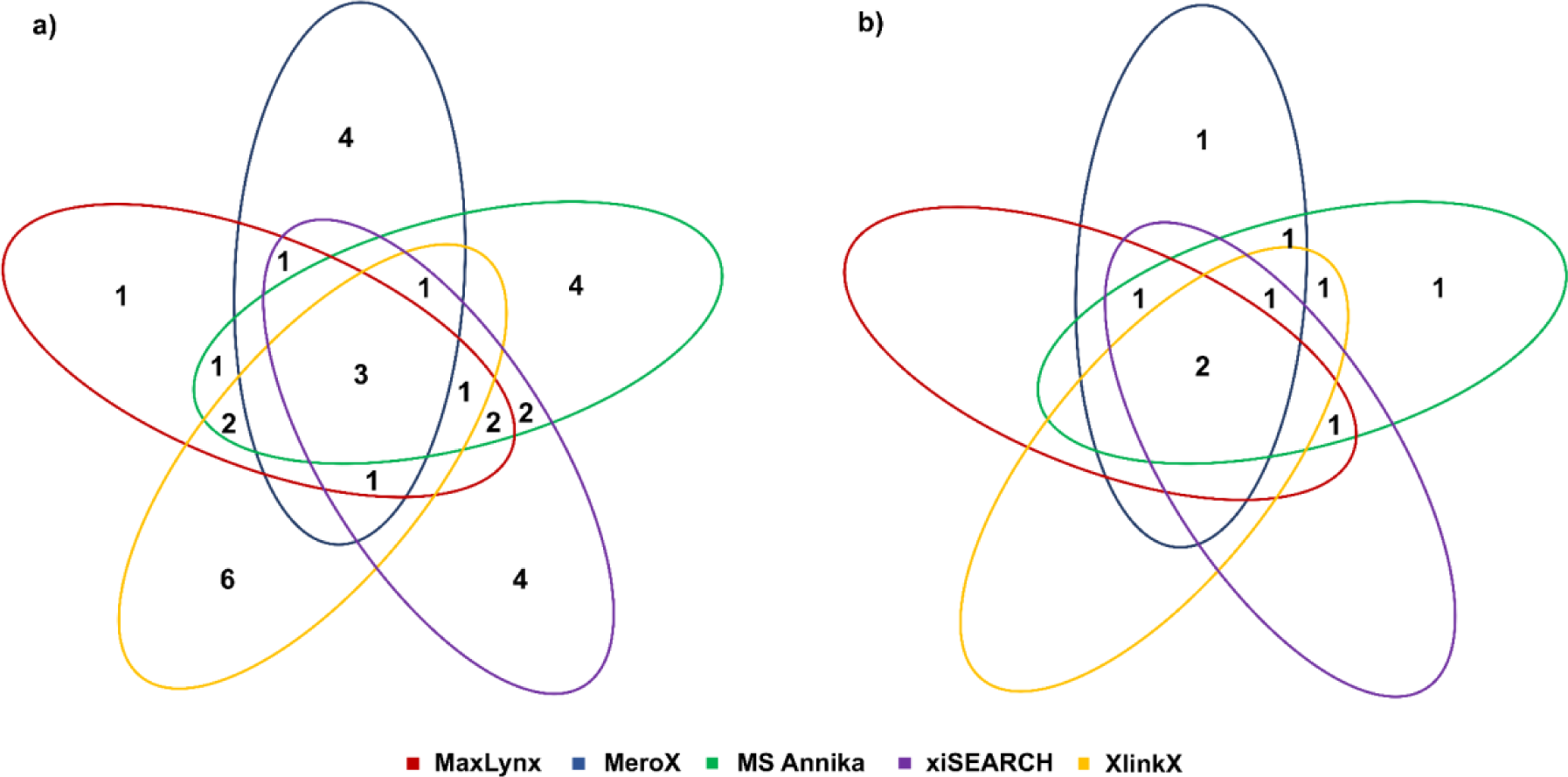
Venn diagram of inter-protein cross-linked residue pairs between complex proteins in a) purified protein sample and b) solubilized membrane sample.

### 3.3. FtsH, HflK, HflC Intra-protein Cross-links

A total of 148 unique intra-protein cross-links were obtained from the two samples. (Supplementary Table 2-3). There were 125, 20, and 3 intra-protein cross-links of FtsH, HflC, and HflK, respectively, from all the software and samples. In the purified protein sample, xiSEARCH gave the highest result with 95 unique cross-links. The lowest number of unique intra-protein cross-links was obtained by XlinkX in the solubilized membrane sample with 24 cross-links. Five of the cross-links were found in all the software considering both samples.

The distribution of intra-protein cross-links was compared in purified protein and solubilized membrane samples separately (Figure 4). There were 10 common intra-protein cross-links obtained from purified protein samples and 13 from solubilized membrane samples with five software. The FtsH-HflK-HflC complex was analyzed with the FASTA files of these three proteins and top 300 proteins from protein search results from MaxQuant (300 proteins and contaminants) in all software. The most abundant 10 inter- and intra-protein cross-links from both samples were listed in Supplementary Table 3. Additional miscleavages and modified peptides were indicated in the table with different symbols. Since serine, threonine and tyrosine modifications were set as modification sides, we clearly observed binding of the cross-linker to a residue other than lysine (Fig. 2).

**Figure 4.**
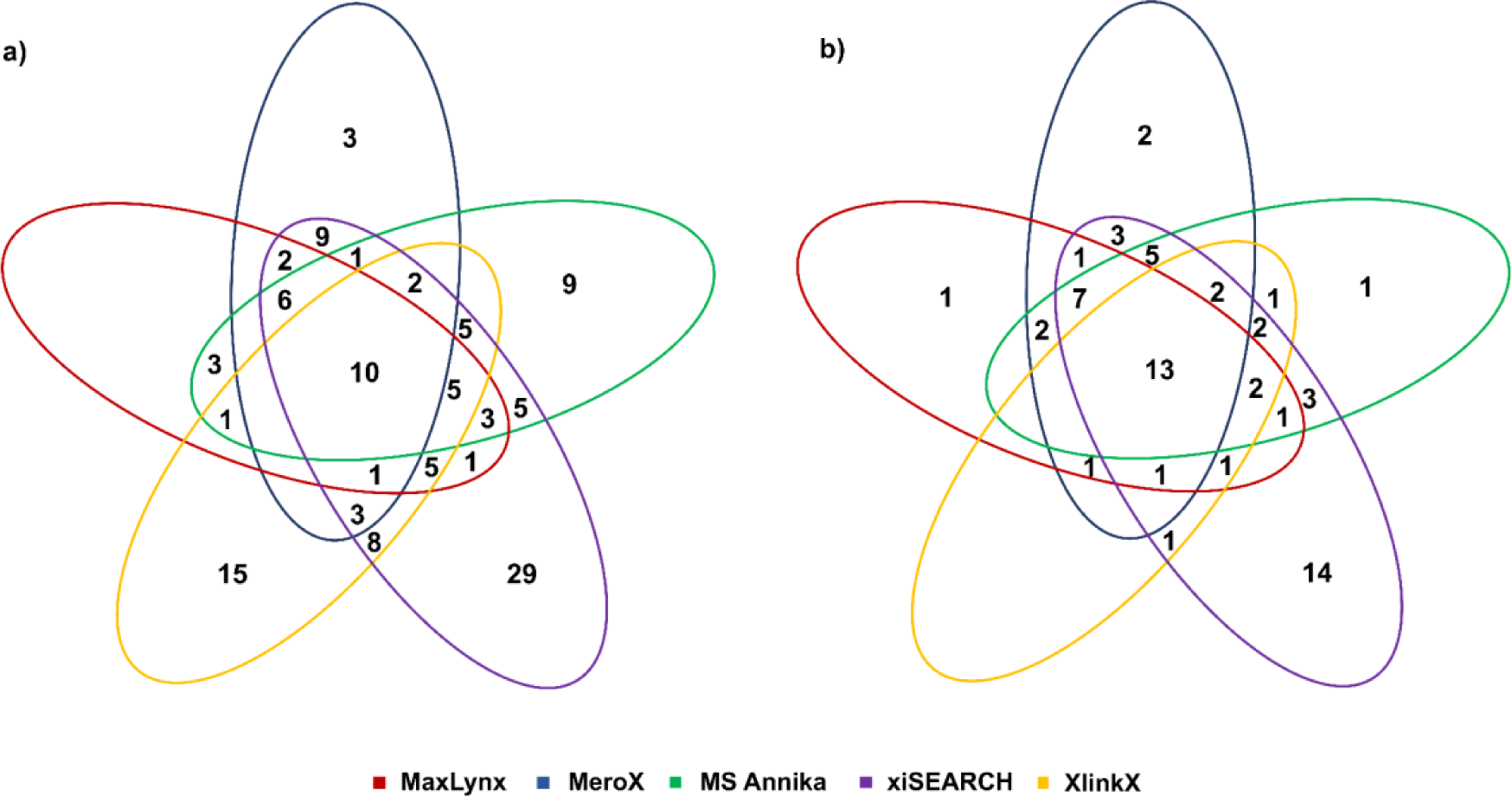
Venn diagram of intra-protein residue pairs within complex proteins a) purified protein sample and b) solubilized membrane sample.

The cross-links were further manually verified using raw data (MS and MS/MS) as an analytical approach although FDR in cross-link search was set to 1%. As suggested in the literature, the actual error could be higher than the targeted error [33]. Therefore, it was aimed to check the cross-linked peptide fragmentation spectra. All of the results obtained from MeroX, MS Annika, xiSEARCH (via xiVIEW [34]) and XlinkX were verified, whereas in MaxLynx, the verification was not performed since the MS/MS spectrum could not be visualized in the software.

### 3.4. Protein Interactions in the Complex

Intra-protein and inter-protein cross-links determined by various XL-MS software were visualized in previously published experimental structures and AlphaFold2-predicted models of the components of the membrane protein complex, FtsH-HflK-HflC in Fig. 5 and Fig. 6. The interaction between HflK and HflC, and their interactions between FtsH were confirmed in the recently published cryo-EM structure of the FtsH-HflK-HflC complex (PDB: 7WI3) [21]. In the cryo-EM structure, HflK and HflC form a heterodimer with a close dimer interface, and the HflK-HflC dimer forms a dodecamer, comprising 12 HflK and HflC monomers. The entire structure of the complex could not be used due to the number of the proteins in the complex and crowded cross-link distribution. A representative model was used to obtain the structure of the entire complex. Thus, the cross-links between HflK and HflC may represent the interactions between either the HflK-HflC heterodimer or neighboring HflK and HflC monomers. Moreover, cross-links between FtsH and HflK-HflC were determined only between the periplasmic region of FtsH, and stomatin/prohibitin/flotillin/HflK/C (SPFH) domain and N-terminal of HflK-HflC proteins, aligning with interactions observed in the cryo-EM structure. Both intra- and inter-protein cross-links obtained by MaxLynx and MeroX were well-distributed on the experimental structure of the membrane protein complex; however, less number of interactions were determined between FtsH and HflK-HflC with MaxLynx and MeroX, compared to those obtained by MS Annika, xiSEARCH, and XlinkX.

**Figure 5.**
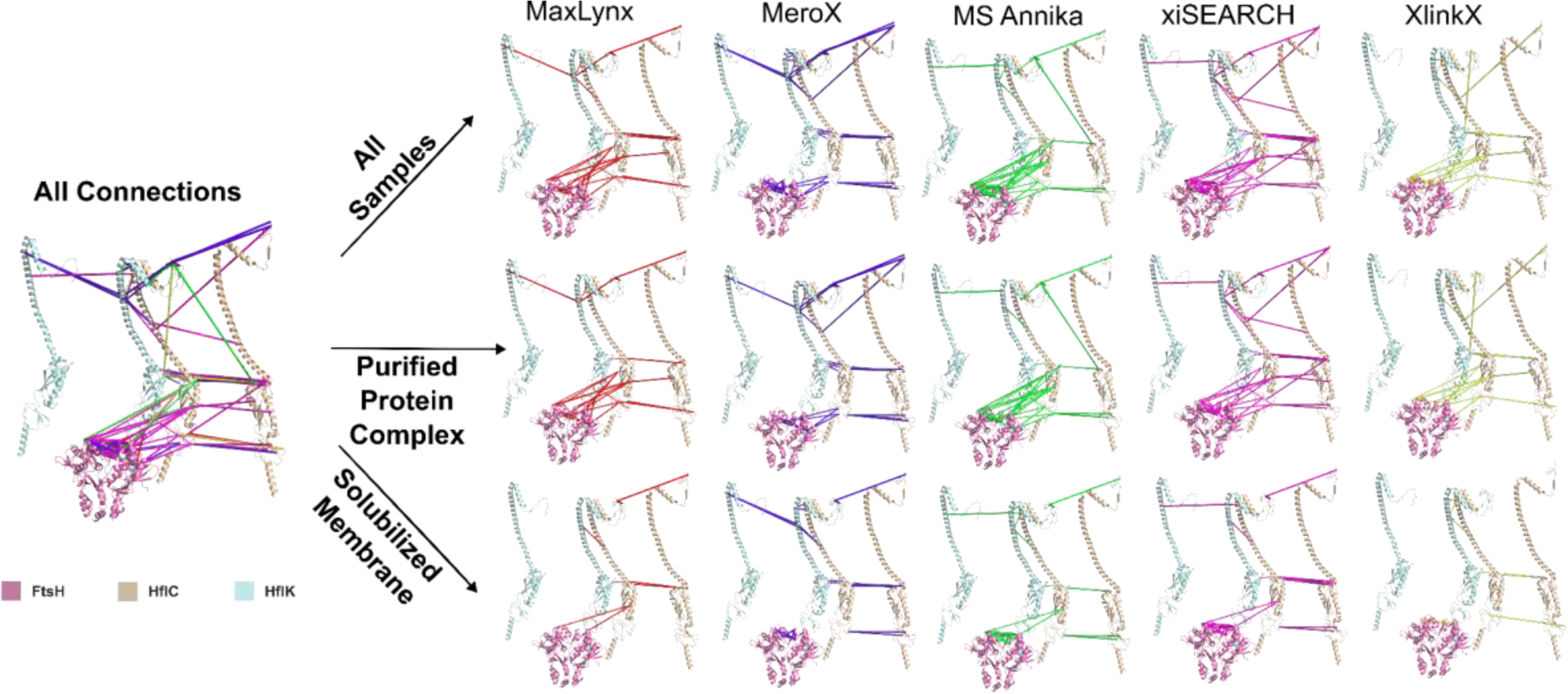
The intra-protein (lines) and inter-protein (dashed lines) interactions of FtsH-HflK-HflC (pink - light blue - wheat) membrane protein complex (PDB ID: 7WI3) obtained from the all XL-MS software and colored as MaxLynx (red), MeroX (blue), MS Annika (green), xiSEARCH (purple) and XlinkX (yellow) in purified protein complex and solubilized membrane samples. Periplasmic and transmembrane regions of hexameric FtsH are shown only. The possible interactions between neighboring HflK and HflC are also shown, represented by the HflK-HflC heterodimers positioned away.

**Figure 6.**
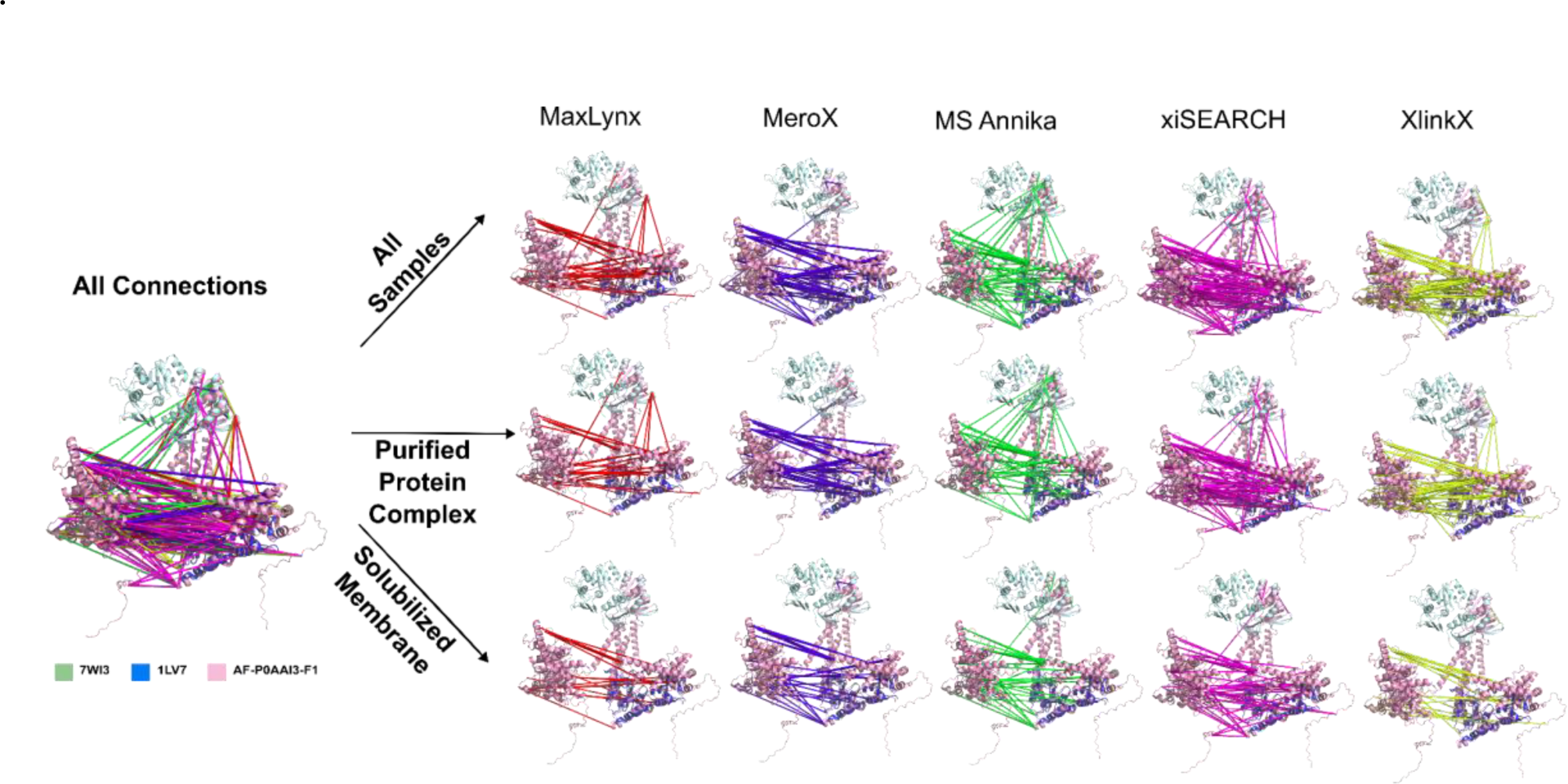
The detailed intra-protein connections of FtsH, represented between two monomers, obtained from XL-MS software MaxLynx (red), MeroX (blue), MS Annika (green), xiSEARCH (purple), and XlinkX (yellow). Experimental structures are superposed with the predicted AlphaFold2 model (pink) (AF-P0AAI3-F1). The periplasmic region of hexameric FtsH (PDB ID: 7WI3) is shown in pale green, the partial cytoplasmic structure (PDB ID: 1LV7) is shown in blue. 91 residues of FtsH, in which no linkage was obtained, in N-terminus were removed in the AlphaFold2 model for clarity.

Two different set-ups result in different cross-link distributions for inter- and intra-protein interactions. The results from both set-ups contain sensible (conformationally possible and generally under 30Å) and some violated (over 30Å and conformationally impossible) interactions. Moreover, the number of cross-linked residue pairs obtained from purified protein samples is higher than that from solubilized membrane samples. Some of the interactions found in the solubilized membrane sample overlap with purified protein sample results. Additionally, certain interactions were only identified in the purified protein sample, not in the solubilized membrane sample, by the same software. The complexity of solubilized membrane in terms of abundant proteins in the system, suppression of the peptide ions during mass spectrometry analysis, and instrumental settings like scan rate might be the reason for this situation. For example, the interaction between FtsH and HflC was not detected in the solubilized membrane sample, but the purified protein complex sample by MeroX and XlinkX. Furthermore, only FtsH and HflK connection was obtained from the purified protein complex sample by MaxLynx.

The intra-protein interactions in FtsH were visualized on both experimental structures (PDB: 1LV7, 7WI3) [21,35] and the predicted AlphaFold2 model (AF-P0AAI3-F1) due to the lack of full-length FtsH structure deposited in the PDB although experimental structures align well with the predicted model (Fig. 6). There are cross-links determined between not only the periplasmic region (N-domain) but also the cytoplasmic region of FtsH. These connections may reflect both intra-protein interactions within the FtsH monomer and between FtsH monomers at the dimer interface as FtsH exists in a hexameric conformation. Purified protein search results contain interactions between the N-domain and cytoplasmic region of FtsH from all five software. However, these interactions appear only in MS Annika and xiSEARCH results of solubilized membrane samples. Moreover, xiSEARCH resulted in a relatively high number of FtsH intra-protein interactions within both the N-domain and cytoplasmic region. On the other hand, there were many interactions determined by MeroX, MS Annika, xiSEARCH and XlinkX within the N-domain; however, there was no cross-link determined in the N-domain with MaxLynx software.

The obtained inter-protein cross-link interactions were located on the full cryo-EM structure of the complex (PDB:7WI3) and the distances between them were measured (Supplementary Table 4). The cross-links were verified whether they were at the suitable distance (30Å) restraint or not. During the evaluation process, two HflK and two HflC as a dimer and neighboring monomers of them with the N-domain of FtsH were used. The shortest distances measured with this model were taken into account. While 65% of the inter-protein cross-links found (24 cross-links) are under the 30Å distance limit, 13% of the inter-protein cross-links found (5 cross-links) are over this limit. 22% of the cross-links (8 cross-links) could not be measured due to the disordered regions in protein structures. The interactions found over 30Å distance might be interactions with other proteins in the medium, as false positives [36].

As for the intra-protein interactions, they were located on the cryo-EM structure of the complex (PDB:7WI3) and cytoplasmic region structure of FtsH (PDB:7WI4). The cross-links were verified by measuring Cα distances with a 30Å distance restraint. The measurement results are listed on Supplementary Table 5. Two HflK and HflC as a dimer and neighboring monomers of them with the N-domain of FtsH were used. Moreover, the cytoplasmic region structure of FtsH was used for FtsH intra-protein interaction evaluation. Since the structure of the FtsH transmembrane region is not available, FtsH intra-protein interactions were checked on two different regions of FtsH cryo-EM structure. According to the shortest distances measured, while 52% of the intra-protein cross-links found (77 cross-links) are under the 30Å distance limit, 14% of the cross-links found to be found (21 cross-links) are over this limit. 34% of the cross-links (51 cross-links) could not be measured due to the lacking regions in the structure.

Structural investigation of two set-ups shows that a small database (consisting of three complex protein sequences) with a purified protein sample gives a higher number of unique residue pairs than a large database (300 protein sequences and contaminant sequences) with a solubilized membrane sample. However, an increasing number of cross-links results in some false positives due to wrong cross-link assignments during the search. On the other hand, results obtained from large database searches give more cross-links in 30Å range and sensible conformations.

Overall, intra- and inter-protein cross-links between FtsH-HflK-HflC were determined by each XL-MS software (Fig. 7a). There are 59 novel cross-links between residues which are not resolved in experimental structures. For instance, HflC-Lys137 interacts with FtsH-Lys138, and HflC-Ser331 interacts with HflK-Lys346 (Fig. 7b). All these cross-links may be used to model the protein complex using experimental and predicted structures, and molecular docking programs such as HADDOCK [37]. Multimeric structures of proteins, for instance hexameric FtsH or dodecameric HflK-HflC, can also be modeled by molecular docking and cross-linking data. Likewise, the multimer structures predicted by structure prediction programs such as AlphaFold [38,39] and RoseTTAFold [40] can be confirmed by XL-MS data. The predicted structures can also be fitted into low resolution cryo-EM maps or SAXS (Small Angle X-ray Scattering) models.

**Figure 7.**
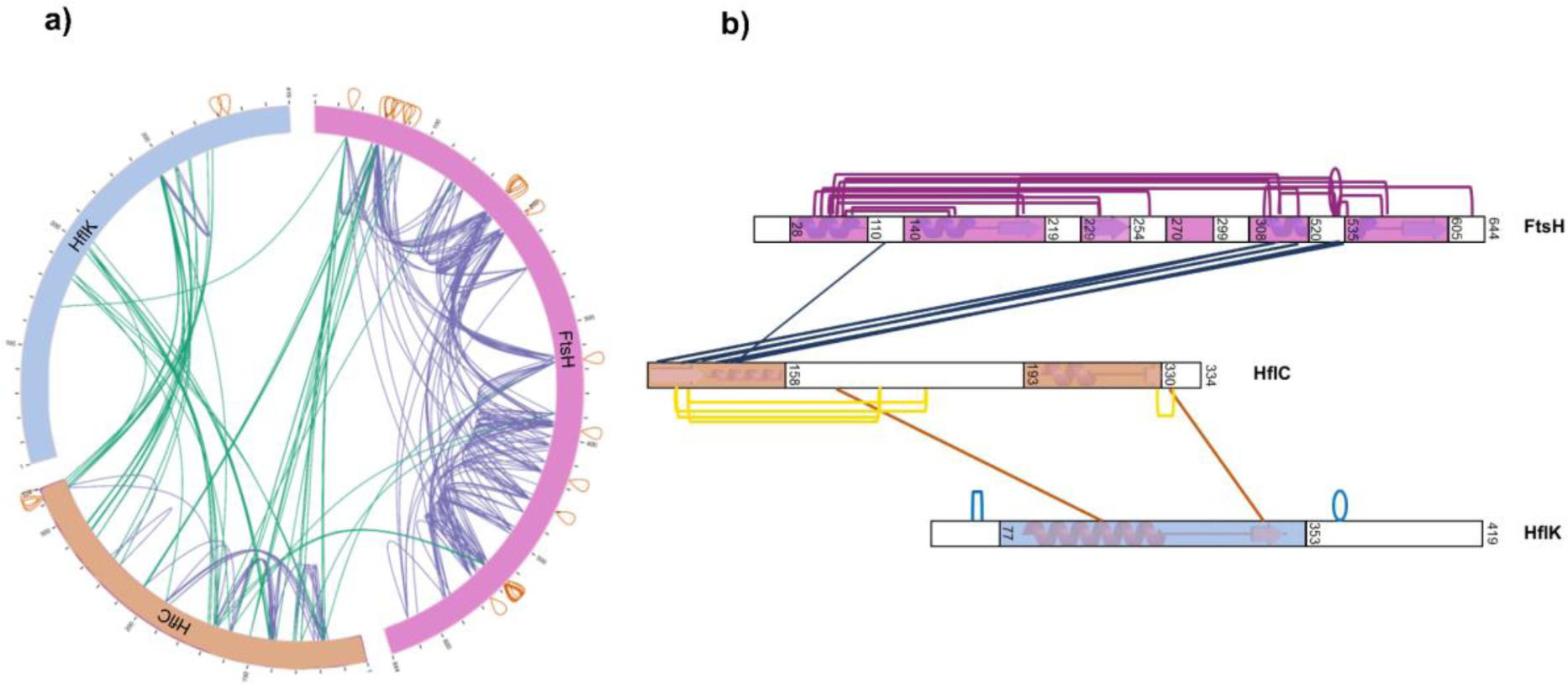
The a) circular network map shows all inter-protein (green), intra-protein (blue), and overlapping (orange) cross-links between HflC (wheat), HflK (blue), and FtsH (pink) proteins in the complex. b) The cross-links corresponding to unstructured regions of complex proteins (PDB: 7WI3, 7WI4) shown on an interaction network map. The unknown regions are colored white, while the known regions are colored with pink (FtsH), wheat (HflC), and blue (HflK). The links between known regions could not be investigated due to the lack of complete complex structure.

In addition to structural proteomics studies, the interaction between proteins can be used for protein mapping and protein-protein networking, especially for complex samples such as a cell membrane as in our study.

### 3.5. XL-MS Software Comparison

Five software were compared according to the number of cross-links identified in two different datasets: solubilized membrane and purified protein complex samples (Figure 8).

**Figure 8.**
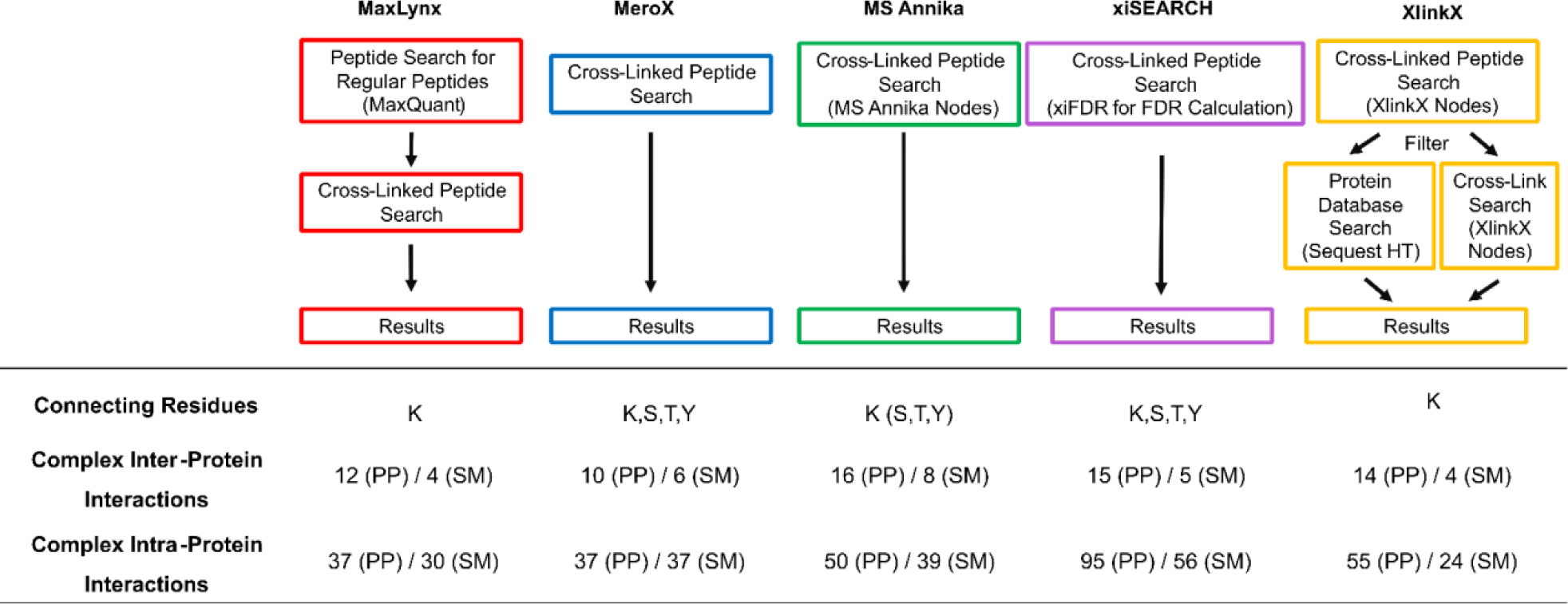
Comparison of five XL-MS software; MaxLynx (red), MeroX (blue), MS Annika (green), xiSEARCH (purple), and XlinkX (yellow). Overall search process, including interacting residues, the number of complex inter-protein cross-links and complex intra-protein cross-links for solubilized membrane (SM) and purified protein (PP) are given.

All the software provided relatively similar numbers of inter-protein residue pair. Considering both samples, MaxLynx provided 16, MeroX did 16, MS Annika did 24, xiSEARCH did 20, and XlinkX did 18 inter-protein cross-link hits. xiSEARCH gave the highest number of intra-protein cross-links in both samples, whereas MaxLynx gave the least.

Proteins were analyzed based on unique peptides. The number of cross-links were based on unique peptides connected via DSBU. The DSBU cross-linker was already integrated into all software. The software outcome file lists inter- and intra-protein cross-links by indicating specific AA positions and corresponding tryptic peptide sequence for each protein pair. The matching score is also listed along with standard MS search parameters such as number of matching spectra, precursor, mass error, retention time and intensities.

MaxLynx is a software embedded into MaxQuant, a protein database search software, and thus, the cross-link search is performed only after the protein search. It is a useful process to identify proteins and their interactions simultaneously.

The cross-linker connection side could be set to only lysine residue by default in MaxLynx, contrary to MeroX and xiSEARCH software which conduct only cross-linked peptide search at a time. Unlike MaxLynx, MS Annika and XlinkX, MeroX and xiSEARCH enable selecting multiple residues such as serine, threonine, and tyrosine in addition to lysine simultaneously. In addition, the graphical user interface (GUI) is user-friendly and fluent, and the size of data is not a limitation. However, in MeroX, the protein search is not an option, and only cross-linking information can be obtained.

MS Annika is a Proteome Discoverer 2.5 node, and the workflow used for MS Annika contains a spectrum selector mode. MS Annika allows conduct searches with multiple residues such as serine, threonine, and tyrosine in addition to lysine, but not simultaneously. It also contains both protein IDs and cross-link information.

xiSEARCH is a software for the identification of cross-linked spectra matches. FDR calculations are carried out with xiFDR software after cross-link search via xiSEARCH. xiSEARCH does not have a spectrum view option but web service xiVIEW provides a spectrum viewer. However, it is important to consider that an increase in file size and the number of protein sequences in the FASTA format exerts a significant impact on the search space. Consequently, increasing the number of proteins poses challenges to complete the search efficiently.

XlinkX is another Proteome Discoverer 2.5 node with a workflow consisting of different nodes, such as Sequest HT. Thus, the protein database search is performed with the cross-link search. Although it is useful to obtain protein search data, the cross-link search performed with only lysine residue is one of the limitations of XlinkX.

## 4. Conclusion

In this study, we demonstrated that XL-MS is a powerful technique to elucidate the protein interactions within a mega membrane protein complex containing FtsH, HflK, and HflC, over-expressed in *E. coli,* using software tools, MaxLynx, MeroX, MS Annika, xiSEARCH and XlinkX. Among five software, MS Annika gave the highest number of inter-protein cross-links, whereas most of the intra-protein cross-links were obtained from xiSEARCH. A higher number of the FtsH-HflK-HflC inter- and intra-protein cross-links were obtained from the purified protein complex sample than the solubilized membrane sample. The findings indicate that simple systems like purified protein samples exhibit greater reliability and yield more cross-links compared to complex systems like solubilized membrane samples. It’s important to note that each software employs a distinct method for calculating the False Discovery Rate (FDR), which influences the efficacy of false discovery control. Therefore, it was necessary for this study to manually confirm the cross-links on the raw MS and MS/MS data to approach the data from an analytical perspective.

The outcomes of the study were assessed onthe complex structure obtained from Protein Data Bank (PDB) to validate intra- and inter-protein cross-links on solved structure (PDB:7WI3). Furthermore, cross-links between residues not resolved in experimental structures were determined. These cross-links may be used to model monomeric and multimeric protein complexes using experimental and predicted structures, and molecular docking programs, assisted with protein structure/assembly prediction programs. Outcomes provided valuable cross-link information within the over-expressed FtsH-HflK-HflC complex. XL-MS can be used for protein mapping and protein-protein networking studies in various matrices. Further studies are ongoing to elucidate the biological phenomena behind the protein-protein interactions.

## Data Availability

Data will be made available on request.

## Supplementary Material Description

Supplementary Table 1. Cross-link search parameters and search database

Supplementary Table 2. Total interaction numbers and connections represented

Supplementary Table 3. The most abundant 10 inter- and intra-protein cross-links

Supplementary Table 4. Inter-protein connection distances

Supplementary Table 5. Intra-protein connection distances

## ACKNOWLEDGMENT

H.A., H.K.I., G.G., and B.V.K. are supported by TUBITAK (Turkish Scientific and Technological Research Council) BIDEB 2232 Program (Project No: 118C225). We thank Lebrilla Group from University of California, Davis, USA for supporting us in cross-linking mass spectrometry analyses. Also, we thank Bergers Lab - Prof. Christiane Schaffitzel-Berger and Prof. Imre Berger from University of Bristol, UK for providing the plasmid expressing the membrane protein complex. Lastly, we acknowledge Dr. Andrea Graziadei for his critical reading of the manuscript and recommendations.

